# Orbitofrontal neuron ensembles contribute to inhibitory control

**DOI:** 10.1101/452938

**Authors:** Pragathi Priyadharsini Balasubramani, Benjamin Y. Hayden

**Affiliations:** Department of Psychiatry University of California San Diego La Jolla, CA. 92037; Neuroscience and Center for Magnetic Resonance Research University of Minnesota Minneapolis, MN. 55455

## Abstract

Stopping, or inhibition, is a form of self-control that is a core part of adaptive behavior. We hypothesize that inhibition commands originate, in part, from the orbitofrontal cortex (OFC). We recorded activity of OFC neurons in macaques performing a stop signal task. Decoding analyses revealed a clear difference in ensemble responses that distinguish successful from failed inhibition that begins after the stop signal and before the stop signal reaction time. We also found a different and unrelated ensemble pattern that distinguishes successful from failed stopping before the beginning of the trial. These signals were distinct from, and orthogonal to, value encoding, which was also observed in these neurons. The timing of the early and late signals was, respectively, consistent with the idea that OFC contributes both proactively and reactively to inhibition. These results support the view, inspired by anatomy, that OFC gathers diverse sensory inputs to compute early-stage executive signals.

## INTRODUCTION

Inhibition is a key function of the brain’s executive control system (Aron et al., 2015; Hampshire and Sharp, 2015; Logan et al., 2015). The process of inhibitory control over a prepotent behavior is often studied with the stop signal task, in which the subject must deliberately withhold a planned action in response to a specific stop-related signal (Eagle et al., 2007; Li et al., 2006; Logan, 1994; Schall, 2001). Performance in this task predicts dysregulated executive function in psychiatric conditions like drug addiction, obsessive compulsive disorder, and obesity (e.g Chamberlain et al., 2006; Iacono et al., 2008; Logan et al., 1997; Nederkoorn et al., 2006; Schachar et al., 1995). One convenient feature of the task is that it is validated in several species, meaning that animal models can be used to provide insight into the mechanisms of human inhibition (Hanes and Carpenter, 1999; Hanes and Schall, 1995; Logan and Irwin, 2000; See Eagle et al., 2008; Pouget et al., 2017, for review). Understanding those mechanisms holds promise in developing rational treatments for psychiatric diseases and may also help address philosophical questions about self-control and the nature of volition (Schall et al., 2002).

In order to stop effectively, our brains must monitor both the sensory world and the internal milieu for information indicating that planned actions have become disadvantageous and should be cancelled. When this control is directly driven by external signals, such as a stop signal, it is known as reactive control (Braver, 2012; Braver et al., 2007; Chen et al., 2010). In the stop signal task (Logan, 1994; Logan and Cowan, 1984), reactive control can be identified because it occurs after the presentation of a stop signal and before the inferred behavioral response to it, the stop signal reaction time (SSRT, See Schall, 2001; Schall et al., 2002 for review). Stopping decisions can also be influenced and in some cases are entirely determined by internal processes, which can begin even before the start of a trial. This form of inhibitory control is known as proactive control (Braver, 2012; Braver et al., 2007; Chen et al., 2010).

Signatures of reactive control have been observed in the frontal eye fields (FEF) and the primary motor cortex, as well as in midbrain structures like the superior colliculus (SC). Many medial prefrontal structures such as supplementary motor area (SMA), pre-SMA, and SEF show signatures of proactive control (Chen et al., 2010; Chikazoe et al., 2009; Majid et al., 2013; Stuphorn et al., 2010; Stuphorn and Emeric, 2012). In the case of eye movements, control is most directly determined by processes occurring in the FEF and SC. In these regions, inhibition is driven by a rapid rise in firing rates of a specific subpopulation of neurons—fixation neurons - that gate the activity of another subpopulation—movement neurons (Hanes and Schall, 1995; Logan et al., 2015; Schall, 1991).

What is the source of these inhibition signals? In our view, inhibition likely does not emerge from a single specialized stopping area, but rather reflects the integration of diverse forms of information, at varying levels of abstraction, bearing on the need to stop (Hampshire and Sharp, 2015). Such pre-inhibition signals are likely to be especially prominent throughout the prefrontal cortex, which, directly and indirectly, is positioned to regulate motor processes (Aron, 2007; Duncan, 2001; Hampshire and Sharp, 2015; MacLeod et al., 2003). We were particularly interested in the orbitofrontal cortex (OFC), a region on the orbital surface that is closely associated with value and reward processing (Padoa-Schioppa, 2011; Rudebeck and Murray, 2014; Schoenbaum et al., 2009; Wallis, 2007). The OFC is the major input for sensory information into the PFC: it receives strong visual, auditory, gustatory, and olfactory inputs. It also has access to signals relating to internal states, via the medial network (Öngür and Price, 2000). It is also often proposed to occupy an early position in PFC processing hierarchies (Carmichael and Price, 1994, 1996; Fuster, 1988; Fuster, 2001; Rushworth et al., 2012; Rushworth et al., 2011). These facts raise the possibility that OFC serves as a first stage (or at least a relatively early stage) for computing preliminary executive signals including, potentially, inhibition ones.

The contribution of the OFC to inhibition remains disputed. On one hand, multiple studies give the OFC a prominent role in inhibition (Bryden and Roesch, 2015; Chikazoe et al., 2009; Dias et al., 1996; Eagle et al., 2007; Horn et al., 2003; Majid et al., 2013; Mishkin, 1964; Roberts and Wallis, 2000). On the other, a good deal of work argues against such a direct inhibitory role of OFC (Chudasama et al., 2006; Ghods-Sharifi et al., 2008; Rudebeck and Murray, 2014; Schoenbaum et al., 2003). Complementary work suggests that inhibition may be the specific domain of functionally and anatomically distinct structures, especially ones located on the ventrolateral surface (Aron et al., 2015; Aron et al., 2003; Banich and Depue, 2015). Finally, another current of research sees the function of the OFC as purely or primarily economic, and thus unlikely to play any major role in inhibition per se (Padoa-Schioppa, 2011; Wallis, 2007). However, recent work indicates that the OFC may have a clear executive role above and beyond its economic one (Bryden and Roesch, 2015; Mansouri et al., 2014; Mansouri et al., 2009; Sleezer et al., 2016; Sleezer et al., 2017). This executive role may include a contribution to inhibition among other functions. Supporting this hypothesis, a recent study found greater spatial selectivity in rodent OFC on successful inhibition trials than on failed inhibition trials, indicating that OFC carries executive signals that directly regulate inhibitory processes (Bryden and Roesch, 2015).

We hypothesized that OFC participates in stopping decisions by modulating responses in downstream regions that in turn regulate motor activity. In an oculomotor task, such signals would have to have appropriate timing to influence stopping. Thus, we predict that successful vs. failed stopping would be correlated with different neuronal ensemble patterns with specific timecourses. We therefore examined ensemble states of OFC neurons recorded in a stop signal task. We found a significant coding pattern difference that emerged following the stop signal but before the stop signal reaction time. We also found a distinct (i.e. statistically orthogonal) ensemble pattern difference that was observable before the trial onset and that derived from the same set of neurons. Both of these patterns were distinct from (also orthogonal to) economic (i.e. value) signals, which were also carried by the same neurons. Together, these pattern differences provide evidence that OFC carries information sufficient to influence inhibition and suggest it may do so both reactively and proactively.

## METHODS

### Subjects

Two male rhesus macaques (Macaca mulatta, subject J and subject T) served as subjects. All animal procedures were approved by the University Committee on Animal Resources at the University of Rochester and were designed and conducted in compliance with the Public Health Service’s Guide for the Care and Use of Animals.

### Recording site

A Cilux recording chamber (Crist Instruments) was placed over the area 13 of OFC (*Figure 1B*). The targeted area expands along the coronal planes situated between 28.65 and 33.60 mm rostral to the interaural plane with varying depth. Position was verified by magnetic resonance imaging with the aid of a Brainsight system (Rogue Research Inc). Neuroimaging was performed at the Rochester Center for Brain Imaging, on a Siemens 3T MAGNETOM Trio Tim using 0.5 mm voxels. We confirmed recording locations by listening for characteristic sounds of white and grey matter during recording, which in all cases matched the loci indicated by the Brainsight system.

**Figure 1.**
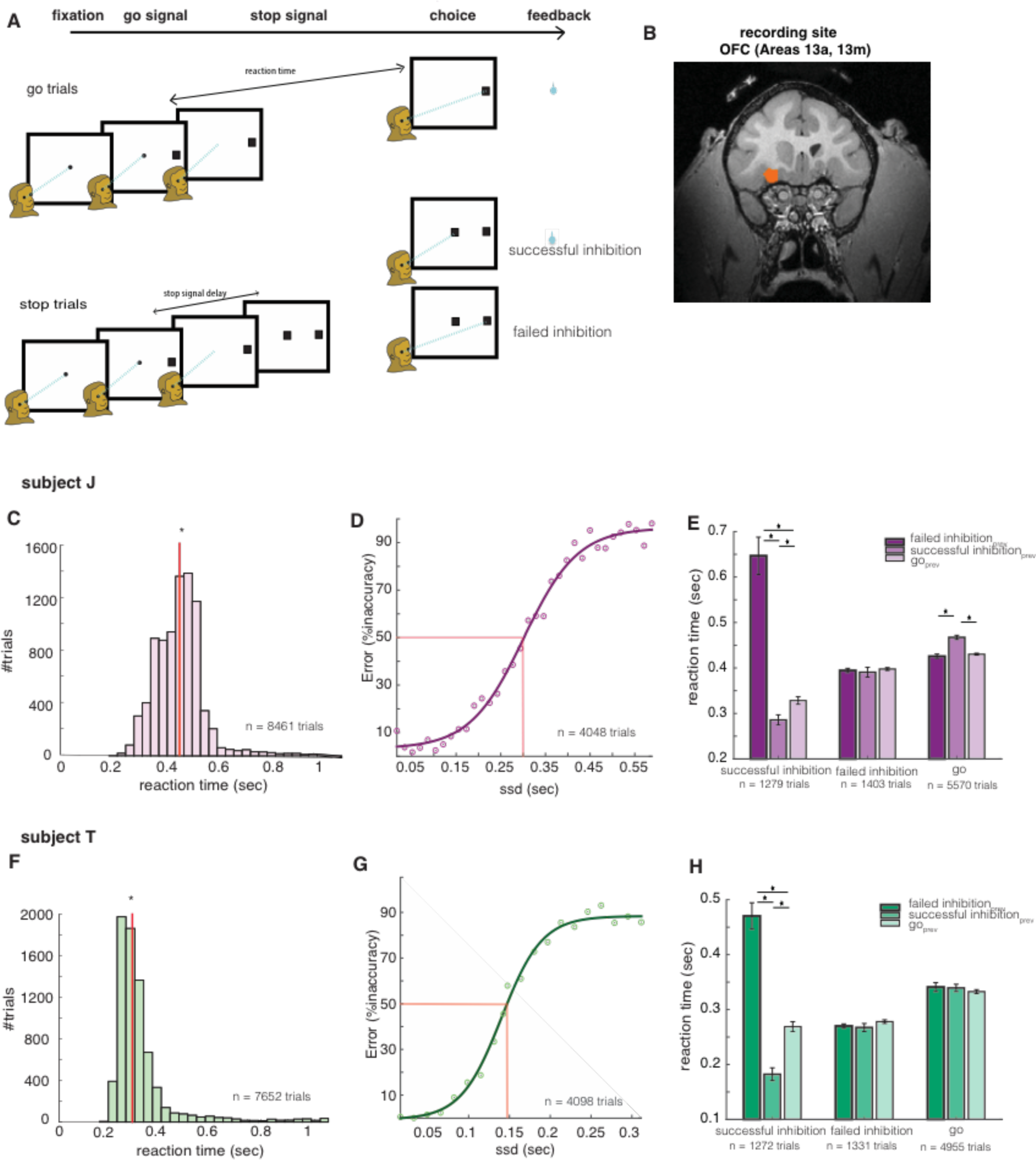
stop signal task and subject behavior: **(a)** task framework **(b)** recording site - Area 13 of OFC (scan from subject J shown) Behavioral results for subject J are presented in panels (c, d, e) and for subject T in (f, g, h) **(c, f)** Go-trial reaction time distributions **(d, g)** Inhibition function varied as a function of SSDs **(e, h)** Previous trial had effects in reaction time behavior. Red lines in (c, e) denote the summation SSD-50_+_ SSRT (Logan 1984; Logan and Cowan, 1984), and red-lines in (d, g) denote SSD-50. Error bars in (e, h) represent SEM, and * denotes t-test significance with p < 0.05.

### Electrophysiological techniques

Single electrodes (Frederick Haer & Co., impedance range 0.8–4 MOhm) were lowered using a microdrive (NAN Instruments) until waveforms of between one and five neuron(s) were isolated. Individual action potentials were isolated on a Plexon system. Neurons were selected for study solely based on the quality of isolation; we never preselected based on task-related response properties.

### Eye tracking and reward delivery

Eye position was sampled at 1,000 Hz by an infrared eye-monitoring camera system (SR Research). Stimuli were controlled by a computer running MATLAB (Mathworks) with Psychtoolbox (Brainard and Vision, 1997) and Eyelink Toolbox (Cornelissen et al., 2002). A standard solenoid valve controlled the duration of water delivery. The relationship between solenoid open time and water volume was established and confirmed before, during, and after recording.

### Task paradigm

The task followed standard stop signal task paradigm (Logan, 1994; Logan and Cowan, 1984). Subjects were placed in front of a computer monitor (1920x1080px) with black background. Following a brief (300 msec) central fixation on a white circle (radius 25px, *Figure 1*), the fixation spot disappeared on the appearance of eccentric saccade target (90px white square, 2.38 degrees, positioned at 288px in left or 1632px in right of screen, 50% chance). A go trial (67% of trials, randomly selected) was indicated by a go signal which is the peripheral target, whereas a stop trial (33% of trials, randomly selected) was indicated by an additional appearance of a stop signal—a central gray square (90px square, 2.38 degrees) delayed relative to the go signal presentation. Stop signal delays (SSD) in the task were set to stabilize at a delay causing approximately 50% successful stopping out of all stop trials recorded for the task in that day; SSDs were modulated through a staircase procedure with intervals of 16 msec. On go trials, subjects were rewarded for a saccade to the go signal and fixating on it for 200 msec; and on stop trials, subjects were rewarded for inhibiting their saccade to go signal and fixating at the stop signal for 400 msec. Water rewards were provided as feedback, and they were contingent on subject’s performance. Rewards were always 125 μl. The inter trial interval was 800 msec.

The *economic choice task* had a similar task framework to stop signal task, and they interleaved randomly in an interval of 1-3 trials. In go trials (random 67% of the total), a peripheral target called go offer (90px white square, 2.38 degrees, positioned at 288px in left or 1632px in right of the screen, 50% chance) was presented, and it was randomly associated with low (15μl), medium (125μl), or high (250μl) reward offers, as indicated by yellow, blue and magenta colored squares, respectively. In stop trials (random 33% of the total), a center stop offer (90px square, 2.38 degrees) delayed with respect to the appearance of go offer was presented in addition. The stop-offer was also randomly associated with yellow, blue and magenta colors to indicate low, medium and high reward sizes. The go offer in stop trials was always in blue color to represent medium reward sized offer. This setup allowed the subject to make a choice through reward comparison in case of stop trials, and through a forced choice in case of go trials. All other parameters were the same as stop signal task.

### Behavioral analysis

Inhibition function related failed inhibitions to stop signal delay (SSD). The delay from the presentation of go signal that caused 50% successful cancellation in stop signal task (SSD-50) was used for computing stop signal reaction time (SSRT). SSRT was usually computed through median and integration methods (Logan, 1994; Logan and Cowan, 1984; Verbruggen and Logan, 2008). *Median* method computed median of go trials’ reaction time distribution and then subtracted SSD-50 from it to give SSRT. The *integration* method computed the point in go trials’ RT distribution whose area was half the whole and then subtracted SSD-50 from it to give SSRT. SSRT computed from both of the above methods gave nearly equal results, and they were averaged to obtain the final SSRT estimates reported for both subjects.

### Statistical methods

Separate PSTH matrices were constructed by aligning spike rasters to the presentation of the go signal and stop signal for every neuron. Firing rates were calculated in 1 msec bins but were generally analysed in longer epochs. Normalization procedure was carried out by subtracting the mean firing during inter-trial interval (ITI) time period and then by zscoring each neuron’s data. For display, PSTHs were smoothed using 200 msec running boxcars. Tests used in the study include two sample t-test for parameteric analysis, Wilcoxin rank test for non-parametric analysis, chi-square test for comparing decoder’s classification accuracy against baseline (50% classification accuracy), Pearson correlation method for correlation analysis. To compute population tuning, we picked neurons with significant (p < 0.05) differences between successful and failed inhibition trials using Wilcoxin rank test.

### Decoding analyses

We chose a neural network based decoding technique because it could efficiently analyse population responses in frontal cortex that are highly multiplexed and non-linear. To generate population activation states as input patterns for the decoding analysis, we first separated all trials of each neuron by trial conditions (successful and failed inhibition trials). Then, we averaged the activity from randomly sampled 10 trials belonging to a condition, with replacement, to form activation state for a neuron in any particular time period. The averaged responses of all 96 neurons’ were pooled to generate one population activation state for a particular trial condition and for a specific time period. 100 unique activation patterns were used for the network training. The procedure is similar to that carried out by other studies (e.g., Mante et al., 2013; Pouget et al., 2000; Rigotti et al., 2013; Wang and Hayden, 2017)

The network used to study the stopping patterns had 100 hidden nodes, and 2 output nodes each representing one target condition for classification. The number of input nodes equal to the total number of neurons used for analysis = 96 (from two subjects). The network weights were initialized to small random numbers between −0.01 and 0.01.

The following *back-propagation algorithm* was used for training the decoders (Haykin and Network, 2004; Rumelhart et al., 1986; Werbos, 1974). In the below, the input nodes are denoted by subscript, *k*, hidden nodes by subscript, *j,* and output nodes by subscript, *i*. Output error, *e*, associated with the network’s response for the *p*’th input pattern was

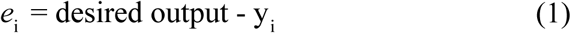

where *y*_i_ was the *i*’th output node response, and desired output was 1 / 0 if the *i*’th output node was associated with target trial condition for the corresponding input pattern (e.g., successful inhibition, failed inhibition). Total output error over all input patterns was computed by,

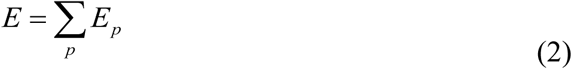

Network’s objective was to minimize the squared output error (eqn. 1) for the *p*’th pattern as denoted by eqn. (3).

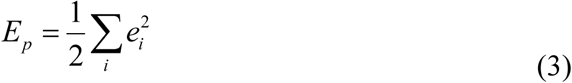

Response of any node was a hyperbolic tangent function (*g*) of slope = 5 of the total input 
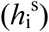
 to it. The output node response, *y*_i_, as a function of its input was calculated as,

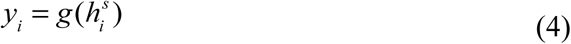

where, net input 
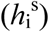
 to the output layer was,

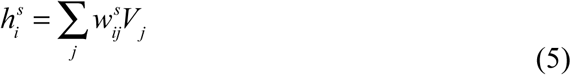

In the above, the weights, *w*_ij_, with superscript, *s*, indicate the second level of the network between hidden and output layer. *V*_j_ denoted the output of hidden layer, and it was represented as a function of net input to the hidden node 
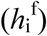
 as follows,

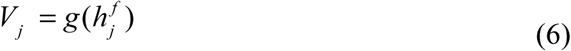

and

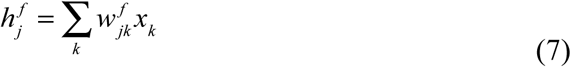

The superscript, *f*, in eqns. (6, 7) denote first level of the network between input and hidden layer, *w*_jk_ were their weights, and *x*_k_ was the input pattern to neural network. Weight updates were proportional to the negative change in error for the *p*’th pattern, *E_p_*, on change in weights. All updates happened trial by trial in the training phase. The update used at the second level was by eqn. (8), and that in the first level was by eqn. (10).

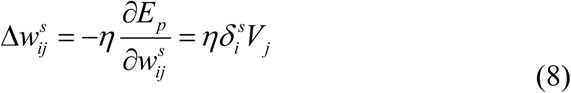

where,

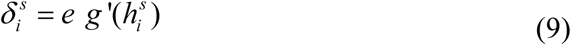

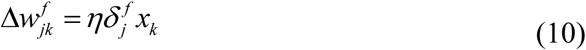

where,

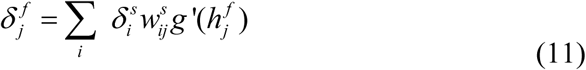

*η* is the learning rate set to 0.001 for pre-go and post-stop signal decoder, and 0.01 for reaction time decoder, and *g*’ denotes first order derivative of hyperbolic tangent function.

We had two different decoders trained on data from 1) pre-go signal, 2) post-stop signal time periods; the former worked on data aligned to presentation of go signal at time = 0, and the latter worked on data aligned to stop signal. For *pre-go decoder,* the training data was population activation states generated on averaging the signal from the fixation epoch spanning 300 msec before the presentation of go signal. For *post-stop decoder*, training data was generated on averaging the firing between 100 msec to 250 msec of stop signal presentation. The entire network was run for *n* = 100 instances with different random weight initializations to obtain average output performance. Training procedure in all instances converged to classification accuracy of above 80%, and the converged weights at the end of training were used for testing of decoder. The testing data used were population activation states generated by averaging 100 msec boxcars that slides with step size of 10 msec (a total of 91 boxcars). The kind of normalization methods didn’t alter the results (See *Figure S2*).

Similarities in the functioning and generalization of pre-go and post-stop decoders were analysed by comparing their converged weights, as well as by comparing their classification accuracy. The similarity index was computed by cross correlating converged hidden layer weight vectors (with zero lag) of two decoders of interest. The index was averaged across *n* (=100) instances of networks with different weight initializations. The similarity index obtained from autocorrelating the weight vectors were used to statistically compare and cross-validate the results from cross correlation, and the results were significant using ttest (ttest, tstat = 210, p < 0.001). Similarities in classification accuracy at pre-go or post-stop signal period were found by using t-test on average performances of the two decoders during *n* instances (with different random weight initializations).

Cancellation time was defined by the size of test-boxcar window positioned at first instance of atleast four consecutive test-boxcars (100 msec window moving in intervals of 10 ms) in a row, whose performance was significantly higher than 50% using chi-square test (p < 0.05). The method avoids false positives that otherwise appear by 99% chance when considering just any one single significant instance of 91 total boxcars. With simulations using markov chains, we found that at least 4 consecutive significant windows were needed in a row for the claim of significance with p < 0.001; so the criteria to find at least 4 consecutive significant bins were used to find pre-go and post-stop decoder results (*figure 4*) as well as cancellation time. Average latency of cancellation signals to SSRT was found by subtracting SSRT of each subject from the mean cancellation time (90 msec).

Incase of the decoder used for analyzing the reaction time ensemble patterns, the inputs to the decoder were either the population activation pattern during time periods 200 msec before or after the reaction time, appropriate to the case of analysis. The output of the decoder was the number of coarse reaction time bins for classifying the input data (n = 5) in the range 0.1 – 0.6 sec. The neurons that contain enough data for all reaction time bins were only considered for this analysis. For reaction time bins taken to be a total of 5, we considered 26 neurons for ensemble analysis. A sum of hundred ensemble patterns was generated for training. The training procedure was similar to pre-go and post-stop signal decoders, and it was carried out for 75% of data. 25% of data was used for testing. The results presented were significant when statistically analyzed against chance percent of 20% using chi-square tests. The decoder results were also cross validated with input as random-data to find that the reaction time ensemble pattern classification was statistically significant with p < 0.001 (ttest, tstat = 27.91).

### Reward and stopping index

Reward index for every neuron was measured by linearly regressing the firing at outcome epoch (between reaction time and feedback) to the received reward sizes in *neuroeconomic trials*. The stopping index was measured as the difference in normalized firing rates (FR) of successful and failed inhibition trials divided by their norm.

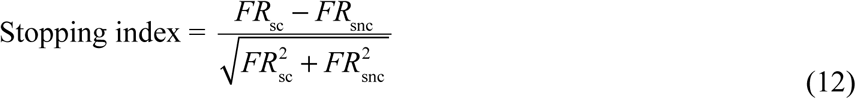

Cross validation tests were performed to support the idea that we had sufficient data to detect an effect had it been there, and to suggest that our results of lack of a significant correlation between stopping and reward indices were statistically meaningful. For the cross validation analysis, all trials within a neuron were randomly separated to two groups, A and B. Stopping and reward index were computed for those two groups of each neuron. We performed correlations between stopping indices of groups A and B, and between reward indices of A and B. A total of *n* (=100) random permutation instances were performed to generate different A and B sets. The test should ideally show high correlations between indices of A and B for any instance, and we indeed saw positive correlations between stopping-index_A_ and stopping-index_B_, and similarly for reward-index_A_ and reward-index_B_. We confirmed that the actual correlation coefficient between stopping and reward indices in OFC fell within bottom 5% of the coefficients computed for *n* instances of stopping-index_A_ and stopping-index_B_. The above was also confirmed for *n* coefficients for reward-index_A_ and reward-index_B_. *Figures 5C-D* showed results of no-significant correlations between stopping and reward indices with p < 0.01.

## RESULTS

### Subjects showed typical behavior in the stop signal task

Subjects performed a standard stop signal task (based on Hanes and Schall, 1995, *Figure 1A* and Methods). On each trial, following a central fixation, monkeys saw an eccentric target (go signal) that, if fixated, provided a juice reward. On a subset of trials (33%, called stop trials), a second signal (stop signal) appeared at fixation and countermanded the previously instructed saccade. Successful inhibition trials were rewarded. Failed trials (trials in which a saccade was made despite a stop signal) were not. Both subjects showed typical behavior in this task; their performance in stop trials varied as a function of time of presentation of stop signals relative to that of go signal (*Figures 1D, 1G*). Median reaction time in go trials was 0.41 sec and 0.27 sec in subject J and subject T, respectively (*Figures 1C, 1F*).

The delay between the go signal and the stop signal is called the stop signal delay (SSD) and it varied randomly across trials. We estimated the SSD that leads to approximately 50% successful stopping (SSD-50) because it can help in computing the stop signal reaction time (SSRT, Logan, 1994; Logan and Cowan, 1984; Verbruggen and Logan, 2008). The SSD-50 was 0.27 sec for subject J and 0.15 sec for subject T. SSRT was 0.14 sec for subject J, and 0.12 sec for subject T. These values are typical of rhesus macaques in these tasks (e.g. Hanes and Schall, 1995; Ito et al., 2003).

Both subjects showed behavioral effects in the reaction times of successful inhibitions as a function of previous trial conditions (*Figure 1E* for subject J*, Figure 1H* for subject T). Successful inhibition trials were shorter when following a successful inhibition trial (subject J: N = 328, subject T: N = 357) as opposed to following a failed inhibition (subject J: N = 111, subject T: N = 60). The statistics for subject J was 360 msec shorter, t-test, t-stat = 11.33, p < 0.0001 and for subject T was 290 msec shorter, t-stat = 11.88, p < 0.0001. Similarly, successful inhibition trials were shorter when following a go trial (subject J: N = 833, subject T: N = 862) as opposed to following a failed inhibition trial (subject J: 310 msec shorter, t-stat = 9.608, p < 0.0001 and subject T: 210 msec shorter, t-stat = 7.72, p < 0.0001).

### Selectivity for stopping in single neurons

We recorded responses of 96 neurons (52 in subject J and 44 in subject T) in area 13 of the OFC (*Figure 1B*). The number of neurons to be collected was determined *a priori* based on exploratory analyses of previous datasets and was not adjusted during recording based on analyses performed mid-experiment. Responses of example neurons are illustrated in *Figure 2*. We focus on neural responses throughout the trial to make it easy to compare the stopping related responses at the time periods before and after SSRT. The responses shown in *Figures 2A and 2B* are aligned to the go signal (time zero). Note that while these response patterns are conveniently illustrative, they do not necessarily stand in for the properties of the entire population (see below).

**Figure 2.**
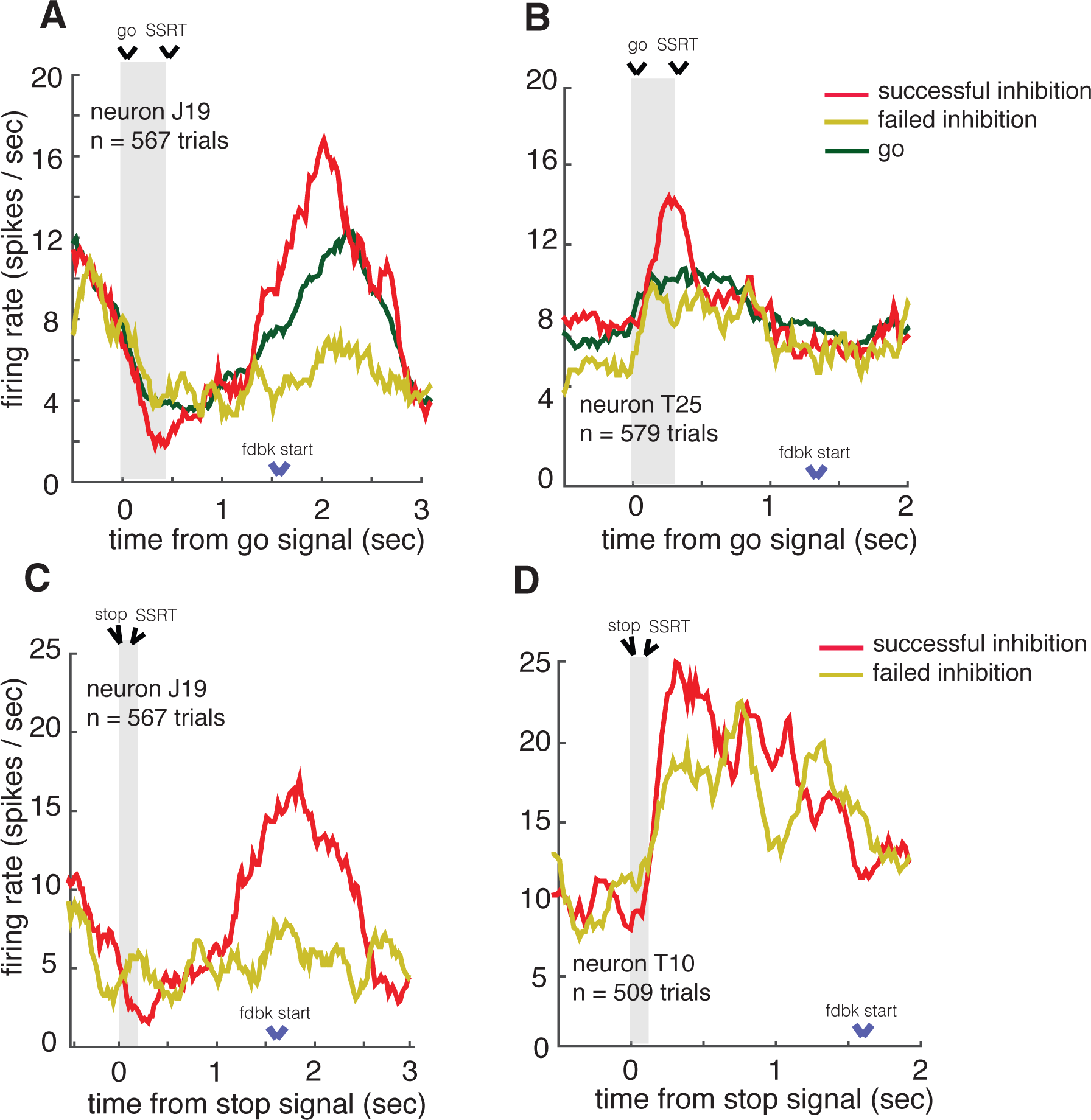
selectivity for stopping in sample neurons: Activity of example neurons during successful inhibition, failed inhibition and go trials are illustrated with respect to **(a, b)** go signal **(c, d)** stop signal presentation time. Time from start of the go (stop) signal to SSRT is shaded in panels A and B (C and D). Neuron in panel A shows significant difference in firing rates of successful and failed inhibition trials before SSRT. Neuron in panel B shows difference even before the beginning of trial. Neuron in panel C is the same as panel A, shows significant difference in firing rates of inhibition trials before stopping response time. Likewise, neuron in panel D shows difference around few msecs after SSRT.

In neuron J19 firing rates following the go signal but before the SSRT were lower on successfully inhibited trials (1.8 spikes/sec) than on failed inhibition trials (4.1 spikes/sec, Wilcoxon rank test, ranksum = 1480, p < 0.05, n = 567 trials, *Figure 2A*). Note that there is a larger and more prominent modulation in firing rate later in the trial. Given its timing, this modulation likely relates to outcome monitoring, is too late to influence stopping, and is not of interest here. Another example neuron, T25, showed distinct patterns for successful and failed inhibition trials even 500 msec before the beginning of the trial (ranksum = 2080, p < 0.05, n = 579 trials, *Figure 2B*).

The responses shown in *Figures 2C and 2D* are aligned to stop signal (time zero). *Figure 2C* illustrates the activity of the same neuron shown in *Figure 2A*; its response pattern showed significant differences between successful inhibition trials (1.8 spikes / sec) and failed inhibition trials (4.4 spikes / sec) that begin after the presentation of stop signal but before SSRT (ranksum = 1340, p < 0.05). Finally, neuron T10 (*Figure 2D*) fired more vigorously on successful than on failed inhibition trials at around 100 msec after the SSRT (ranksum = 2229, p < 0.05). Simple population analyses suggest that these individual neurons are somewhat atypical, however.

Relation between stopping and reward related responses for the sample neurons are dealt in detail in the later part of the manuscript.

### Population averages provide weak information about stopping

Our central hypothesis was that OFC predictively distinguishes successful from failed inhibition. We focused our analyses on two time periods of the trial: 1) The post-stop signal, but pre-SSRT period, and 2) the pre-go signal time period. The post-stop epoch is important because it is when inhibition generated in response to countermanding commands would presumably occur, and has therefore been the focus of many studies of stopping (Schall, 2001; Schall et al., 2002; Logan et al., 2015). It corresponds to the time during which reactive control occurs (Stuphorn and Emeric, 2012). The pre-go signal epoch corresponds to a time before the trial begins; signal differences here presumably reflect proactive control (Stuphorn and Emeric, 2012).

Analysis of single neurons did not provide strong evidence for a role for OFC in stopping. The percent of neurons that individually distinguish successful and failed inhibition trials (regardless of sign) was 8.43% during the 100 msec post-stop signal time period, and was 10.50% during the 100 msec pre-go signal time period. (These epochs were selected before analysis in order to reduce the likelihood of p-hacking). These proportions were not significantly greater than chance in either of the two key epochs (chi-square stat = 1.22, p = 0.26 in the post-stop signal time period; chi-square stat = 1.8, p = 0.17 in the pre-go signal time period). This lack of a detectable effect does not imply that a correlation between stopping and unit activity in OFC does not exist; rather it suggests that if it does exist it is too weak to detect using conventional methods that focus on single neurons in a sample of the size we collected.

We next tested whether successful and failed inhibition trials have a consistent sign of effect on firing rates. The percent of significantly positive cells (successful > failed) was 5.40%, and wasn’t significantly different from the percent of significantly negative (successful < failed) cells (3.03% chi-square test, chi-square stat = 0.52, p = 0.47) in the post-stop signal period. The difference in the sizes of the two cell classes was also not significant before the start of trial at the pre-go signal time period (significantly positive cells 7.55%, significantly negative cells 2.95%, chi square = 2.40, p = 0.12).

Next we looked at grand averages of populations of neurons (*Figure 3*). We observed no difference between successful and failed inhibition trials either after the stop signal or before the beginning of trial. Specifically, during the post-stop signal time period, responses were slightly less for successful than failed inhibition in subject J (average of 0.3 spikes/sec, p = 0.6, *Figure 3B*); the opposite pattern was observed in subject T (average of 0.52 spikes/sec, p = 0.53, *Figure 3D*). Neither effect was statistically significant. Thus, these results suggest that conventional population averages don’t reveal information about the pattern of stopping. Together these analyses indicate that, if stopping correlates exist in OFC, they are of a different form than they take in regions like FEF and SC.

**Figure 3.**
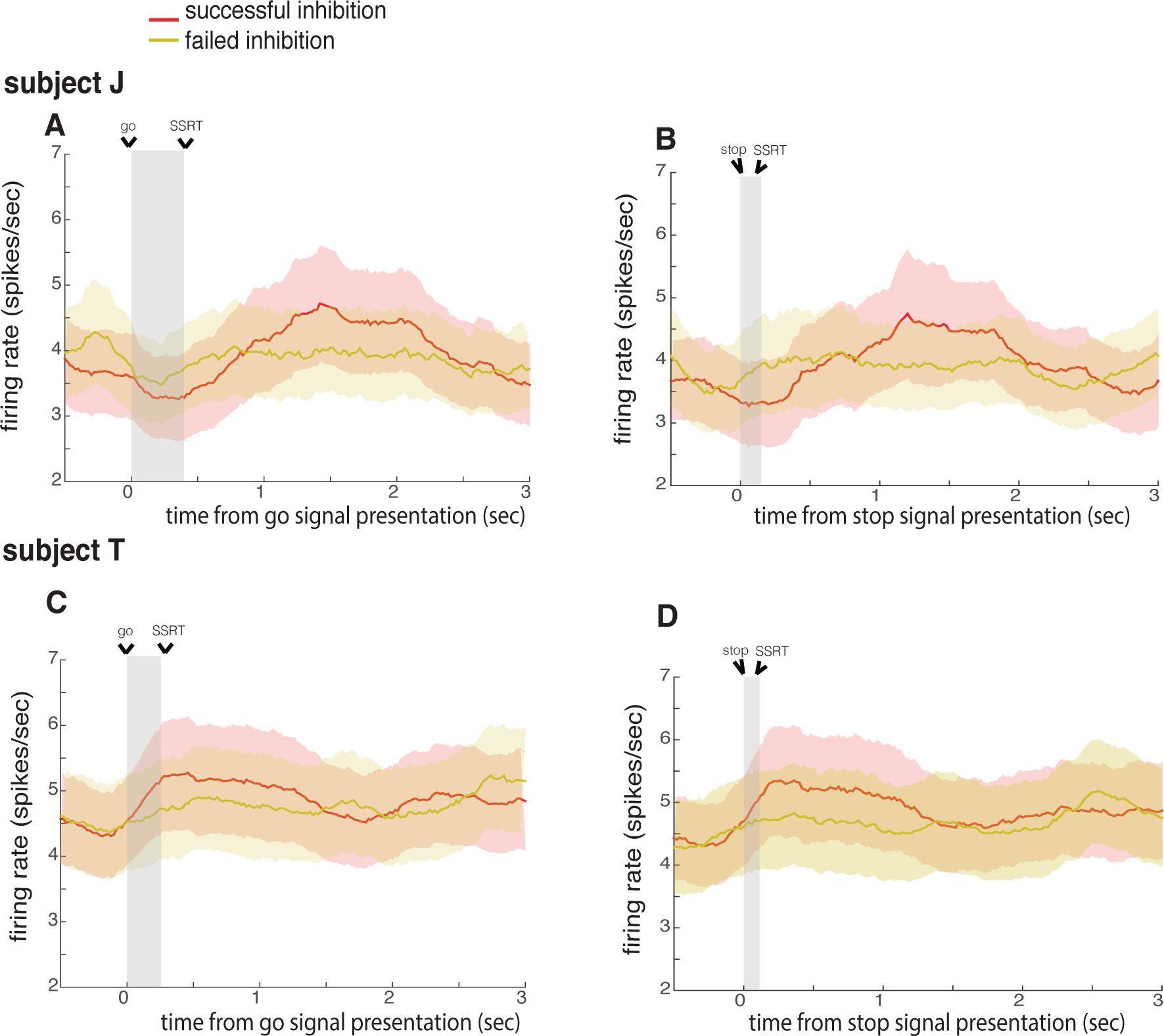
population averages provide weak information about stopping: Population activity for successful inhibition and failed inhibition with respect to **(a, c)** go signal presentation and **(b, d)** stop signal presentation, for subjects J and T. Time from start of the go (stop) signal to SSRT is shaded in panels A and C (B and D). Data for all SSDs are averaged to present successful and failed inhibition trials. Error bars denote SEM. They don’t reveal significant information about the pattern of stopping.

### Ensemble patterns strongly distinguish successful from failed stopping

Many studies suggest that ensemble patterns possess properties to code for neural information and dynamics, which may not be expressed at the level of single units (Averbeck et al., 2006; Meyers et al., 2008; Zemel et al., 1998). Some studies extend these thoughts to suggest that information stored in patterns are more important for neural processing than that present in single units (Morcos et al., 2018). Taking inspiration from such recent developments in the theoretical understanding of neural activity, we devised our next analysis on ensemble patterns. We trained neural network decoders to analyze differences in population activation patterns between successful and failed inhibition that weren’t measured through unit responses or population averages. We were, again, interested in two time periods: 1) the times after the presentation of the stop signal (which we examined using a decoder trained on post-stop signal pattern, referred to below as post-stop decoder), and 2) the time before the start of trial (which we examined using a decoder trained on pre-go signal pattern, referred to below as pre-go decoder). To ensure we had enough data to detect significant effects we used 100 msec moving boxcars and to gain some insight into the time course of effects, we used a 10 msec step size for boxcars.

The post stop signal decoder was able to classify success of an inhibition significantly in a series of 9 consecutive boxcar bins spanning 40 msec after the stop signal to 120 msec after it (these times indicate the starts of the 100 msec boxcars). The numbers for individual subjects were 40 to 140 msec in subject J and 40 to 220 msec in subject T; the central points of the first of these bins is 90 msec for both subjects. These series are unlikely to occur by chance (p < 0.001 in all cases, see Methods for specific use of chi-square tests to quantify significance of consecutive bins, and *Figure S1*). Notably, the central point of the first bin of the series to reach significance in both subjects occurred before the stop signal reaction time of either subjects (the SSRTs were 140 msec for subject J and 120 msec, also see *Figure S1*; *Figure 4B*). We call the central point the cancellation time; it measures the center point latency of first statistically significant difference between successful and failed inhibition trials for the ensemble of neurons. The cancellation time is 90 msec for both subjects. The cancellation time preceded the average stopping response by 50 msec in subject J, and by 30 msec in subject T, suggesting OFC’s responses may precede the stopping response (see Discussion, and *Figure S1*).

**Figure 4.**
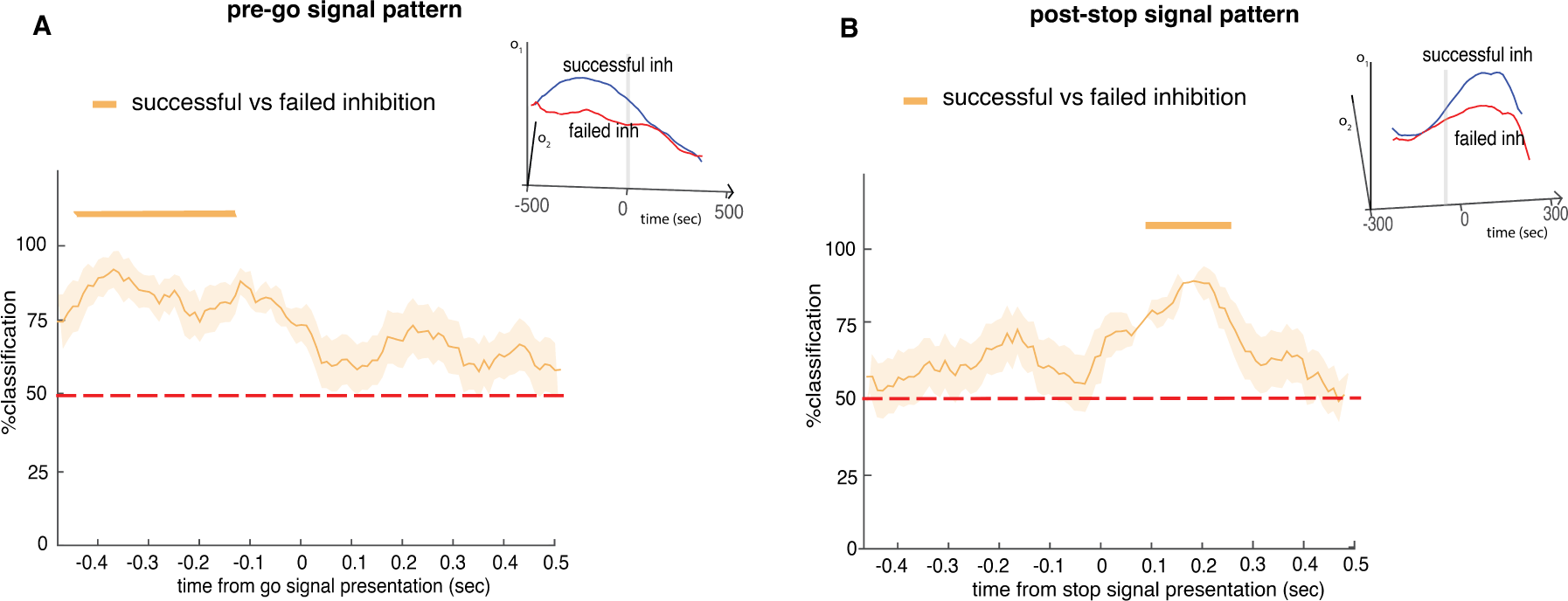
ensemble analysis inform about stopping: Performance of **(a)** pre-go signal **(b)** post-stop decoders to distinguish successful vs. failed inhibition pattern. Insets in both panels illustrate projections of decoder’s output responses (o_1_ and o_2_, for successful and failed inhibition trials, respectively). Error bars represent SEM. Time points highlighted through horizontal bars denote start time of 100 msec boxcars having percent accuracies of classification above chance of 50% (chi-square test, p < 0.05).

We then examined the response differences of the pre-go signal decoder. We observed a significant pattern difference between successful and failed inhibition trials (p < 0.001) extending from 470 to 120 msec before the go signal (see Methods on procedures to determine statistical significance of a boxcar using chi-square statistics). For subject J, significant decoding was observed during the time periods 460 to 120 msec; for subject T it was 420 to 200 msec. These results indicate that the upcoming success or failure of inhibition is decodable from OFC patterns even before the start of the trial (*Figure 4A*, also see *Figure S1*); Our results do not tell us why this correlation exists, although one may infer that it reflects some internal state facilitated by variety of factors such as frequency of task events, frequency of different trial types, motivation, trial sequence, altogether driving successful versus failed inhibition (Chen, Stuphorn, J Neuro, 2010); thus it is a likely correlate of proactive control.

### The post-stop and pre-go decoders are statistically orthogonal

We next examined how the two decoders related to each other. That is, we asked whether the patterns that distinguish successful and failed inhibition after the stop signal related to those that predict inhibition before the trial begins? We did so by comparing the vector of weights of the post-stop decoder and pre-go decoder. We found a very low similarity between them (similarity coefficient, ‘*r*’, obtained at zero lag on cross correlating weight patterns, *r* = 0.02 ± 0.22). This low correlation may be due to noise in our signal. We therefore performed a cross-validation procedure to measure range of values expected from a true correlation with noise. The measured coefficient fell below 1st percentile of that obtained from autocorrelation (and thus is significant at p<=0.01; 100 randomizations, average *r* from the randomized sets = 25.84 ± 0.82). The decoding performances during the time periods after the stop signal (t-stat = 6.0491, p = 0.003) and before the beginning of trial (t-test, t-stat = 8.8874, p < 0.001) were significantly different between the networks trained on pre-go and post-stop signals. The above results suggest that the two decoders that predict successful vs. failed inhibition are statistically orthogonal and thus dissimilar in the early and late epochs.

### Activity of OFC ensembles, but not single neurons, correlates with reaction time

Neurons in prefrontal structures such as SEF show linear correlations between single neural responses and trial reaction time (Stuphorn et al., 2010). We next asked whether single neuron responses from our OFC data around response showed correlations to reaction time, the time taken to saccade to the choice target. We computed reaction time as the time difference between the presentation of go signal and time of saccade to target (on trials with such saccades, go trials and failed inhibition trials, Hanes et al., 1996; Stuphorn et al., 2010).

We used mean firing rates during 200 msec before and after the reaction time for this analysis; our analytical approach was designed to be similar to that used by Stuphorn et al. (2010). We found no correlations between reaction time and firing rates before the reaction time (Pearson correlation, ϱ = 0, p = 0.41). Likewise, we found no correlation after the reaction time (Pearson correlation, ϱ = ‒0.01, p = 0.09). This analysis suggests that activity of single neurons in OFC, unlike those in SEF, doesn’t scale linearly with reaction time (Stuphorn et al., 2010). This lack of observed correlation raises the possibility that downstream regions may not linearly decode the information from OFC for informing the urgency of action execution.

We hypothesized that OFC ensemble responses predict reaction times. To test this idea, we generated population activation patterns from neurons that contained data for discretized reaction time bins in a range of 0.1 sec to 0.6 sec with step size 100 msec (5 equally sized bins, see Methods for the specific criterion used). In particular, we asked whether the ensemble response could be accurately classified to discrete reaction time bins (see Methods) in a non-linear fashion. The results show that neural network decoders were able to classify OFC ensembles to correct reaction time bins (See Methods, ttest, tstat = 27.91, p < 0.001), when the population activation pattern was generated from 200 msec time periods before and after the reaction time (chi-square test, p < 0.05 for all 5 bins). These results suggest that OFC ensemble responses can predict reaction times.

### OFC codes for stopping and reward are unrelated

The reward-encoding role of OFC is a hallmark of its function (Padoa-Schioppa, 2011; Schultz, 2000; Wallis, 2007). We therefore wondered whether the stopping-related activity that we observed might be a side-effect of its reward roles. For example, it may be that there is some undetectable natural variation in the relative subjective value of the reward offered for correct performance. On trials in which the reward happened to have a slightly lower value, the subject would be less motivated to perform correctly; this fluctuation would then introduce a correlation between firing rates and successful inhibition.

To test for the possibility that our putative inhibition signals were just reward correlates, we took advantage of a second set of trials, collected in a *neuroeconomic stopping task*; detailed analysis of the results from that task will be the focus of a later manuscript. In this task, subjects chose or rejected a single reward that had one of three values (low, medium, and high rewards, see Methods). The two task types, neuroeconomic and standard stop signal paradigms, were randomly interleaved on a trial-by-trial basis. The data from this task allowed us to assess each neuron’s tuning function for anticipated rewards. Responses to different reward amounts by two example neurons are shown in *Figures 5A and 5B*. We found tuning for anticipated reward values in the firing activity during the reward feedback time period. For example, we observed a significant positive correlation between reward amount and firing rate in neuron J19 (ϱ = 0.3138, p < 0.001, *Figure 5A, same as Figure 2A but aligned to feedback*) and a significant negative one in neuron T10 (ϱ = ‒0.143, p = 0.04, *Figure 5B*).

**Figure 5.**
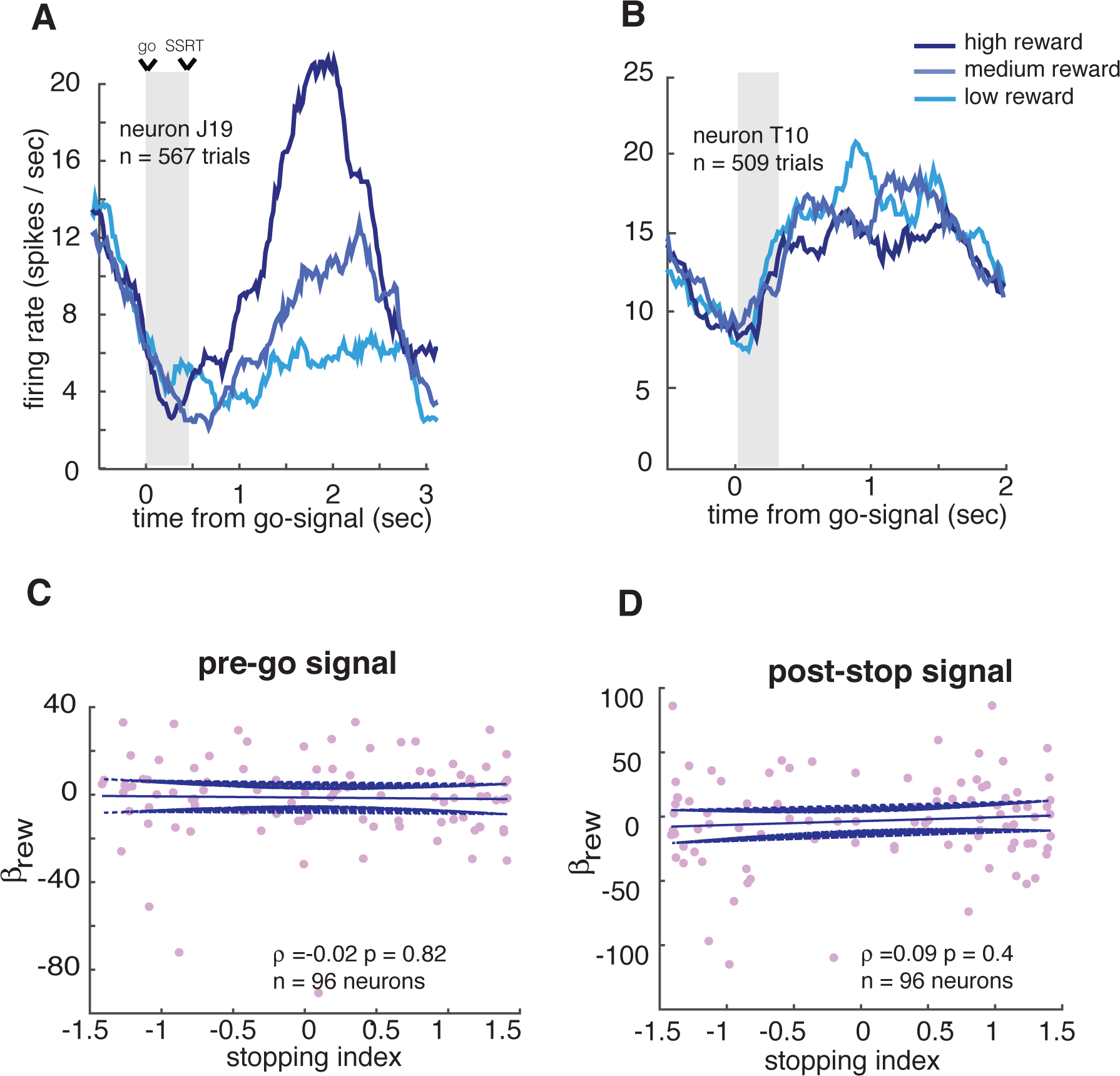
unrelated reward and stopping codes: **(a, b)** Illustration of example neurons tuned to reward sizes. Neuron in panel A (panel B) shows significant positive (negative) correlation to reward amounts. Correlations between stopping and reward indices show no significant effect during 100 msec in **(c)** pre-go signal and **(d)** post-stop signal time period.

If the stopping-related signals were a consequence of reward encoding, we would see a positive correlation between coding patterns for rewards and stopping. We computed a *reward index* for all neurons by regressing their responses to outcomes against the outcomes themselves. We computed a stopping index for all neurons by subtracting on their firing rate during successful and failed inhibition before the stop signal reaction time (see Methods). We found no correlations between these indices in the post-stop signal time period (Pearson correlation, ϱ =0.09, p=0.4, *Figure 5D*). Nor did we find such correlations in pre-go signal time period (ϱ =‒0.02, p=0.82, *Figure 5C*).

This lack of correlation may be a sign that the reward code and the stopping code are different. It may also, in theory, be due to lack of sufficient data to detect a significant effect. To test this idea, we performed a cross-validation analysis (See Methods). Specifically, we reasoned that if insufficient data were the problem then a within sample correlation would also produce no significant correlation. A positive correlation of a within sample correlation, using randomly sampled half-sized subsets, then, would indicate that our data have sufficient power to detect a significant effect (Blanchard et al., 2015). We thus tested whether the correlation coefficient for stopping and reward indices fell below the bottom 5 percentile of the coefficients obtained for within-group correlations. Indeed, the coefficient fell below 1st percentile of that obtained for 100 randomizations in cross validation analysis. *Figures 5C-D* show no-correlations between stopping and reward indices with p <= 0.01.

## DISCUSSION

We examined the correlates of successful stopping in ensembles of neurons in OFC. We found that while traditional single unit correlates of stopping associated with more dorsal structures are too weak to observe (if there at all), that ensembles of neurons in OFC readily distinguish successful and failed inhibition. These signals were not consistently associated with a higher or lower firing rate, nor were they associated with two discrete sets of neurons, as in FEF and SC (Hanes et al., 1998; Pouget et al., 2017; Stuphorn et al., 2000). These findings, then, raise the possibility that OFC plays a somewhat different role in regulation of stopping than these other regions.

Our study shows the presence of stopping-related patterns in OFC at two specific time periods, the first one after the presentation of stop signal, and the second one before the beginning of trial. The timing of the two stopping-related patterns is reminiscent of the times associated with with reactive and proactive control, respectively (Braver, 2012; Braver et al., 2007; Chen et al., 2010; Chikazoe et al., 2009; Hanes et al., 1998; Ito et al., 2003; Majid et al., 2013; Stuphorn et al., 2000; Stuphorn et al., 2010; Stuphorn and Emeric, 2012). These response patterns indicate that OFC neurons carry a signal that precedes stopping decisions. They suggest, then, that OFC may be part of the pathway that determines the success or failure of stopping.

OFC is marked by its unusual anatomy: it receives strong and diverse sensory inputs, as well as visceral ones, and it projects to more dorsal prefrontal structures that collectively directly regulate behavior (Öngür and Price, 2000; Cavada et al., 2000, Wallis, 2007). Its connections then mean that it has a nearly ideal anatomy for monitoring sensory and reward information forming a first draft of the type of executive signals that can inform - but not determine - action. Its influence is unlikely to be limited to inhibition; its executive functions likely include contingent (rule-based) decisions, working memory, switching, and conflict monitoring (Bryden and Roesch, 2015; Lara et al., 2009; Mansouri et al., 2014; Sleezer et al., 2016; Sleezer et al., 2017). More broadly, these executive signals likely constitute a component of a larger set of output prediction signals that determine OFC’s major role in cognition (Rudebeck and Murray, 2014; Schuck et al., 2016; Wilson et al., 2014).

Another well-known role of the OFC is in signaling value (Padoa-Schioppa, 2011; Wallis, 2007). Given this close association, one might expect that its role in inhibition is to signal the current subjective value of stopping. Our data, which indicate that OFC value responses are orthogonal to its stopping responses, argue against this possibility. They are consistent, however, with the broader theory that OFC carries a suite of signals that regulate the ongoing transformation of stimulus into action, and that value is one such signal. Other executive signals observed in OFC include spatial information and abstract rules (Lara et al., 2009; Luk and Wallis, 2013; Sleezer et al., 2016; Strait et al., 2016; Tsujimoto et al., 2009; Wallis, 2007; Wallis et al., 2001), those related to switching and task adjustment (Chase et al., 2012; Sleezer et al., 2017), and to conflict adaptation (Bryden and Roesch, 2015; Mansouri et al., 2014; Mansouri et al., 2009).

In foraging theory, decisions are generally framed as accept-reject. From this perspective, binary choices, the mainstay of behavioral economics and microeconomics, are actually better thought of as two somewhat independent accept-reject decisions (Kacelnik et al., 2011). Each accept-reject decision, in turn, functions like a classic accept-reject foraging decision, that is, as a choice between pursuing and refraining from pursuit. In other words, what appears to be a binary choice may actually be a pair of stopping decisions. If economic choice ultimately boils down to stopping, there is an opportunity for a “grand unified theory” that can explain the two types of decisions. This possibility would help explain, for example, why many of the same regions are involved in both types of decisions. In particular, OFC has demonstrated importance in both economic decisions and in inhibition. Progress in this area promises to help shed light on important debates, such as how economic choice relates to self-control (Berkman et al., 2016; Shenhav, 2017).

## Acknowledgements

This work was supported by an R01 (DA038615) to BYH. We thank Meghan C. Pesce for help with data collection.

The authors declare no conflicts of interest.

## SUPPLEMENTARY MATERIAL

**Figure S1.**
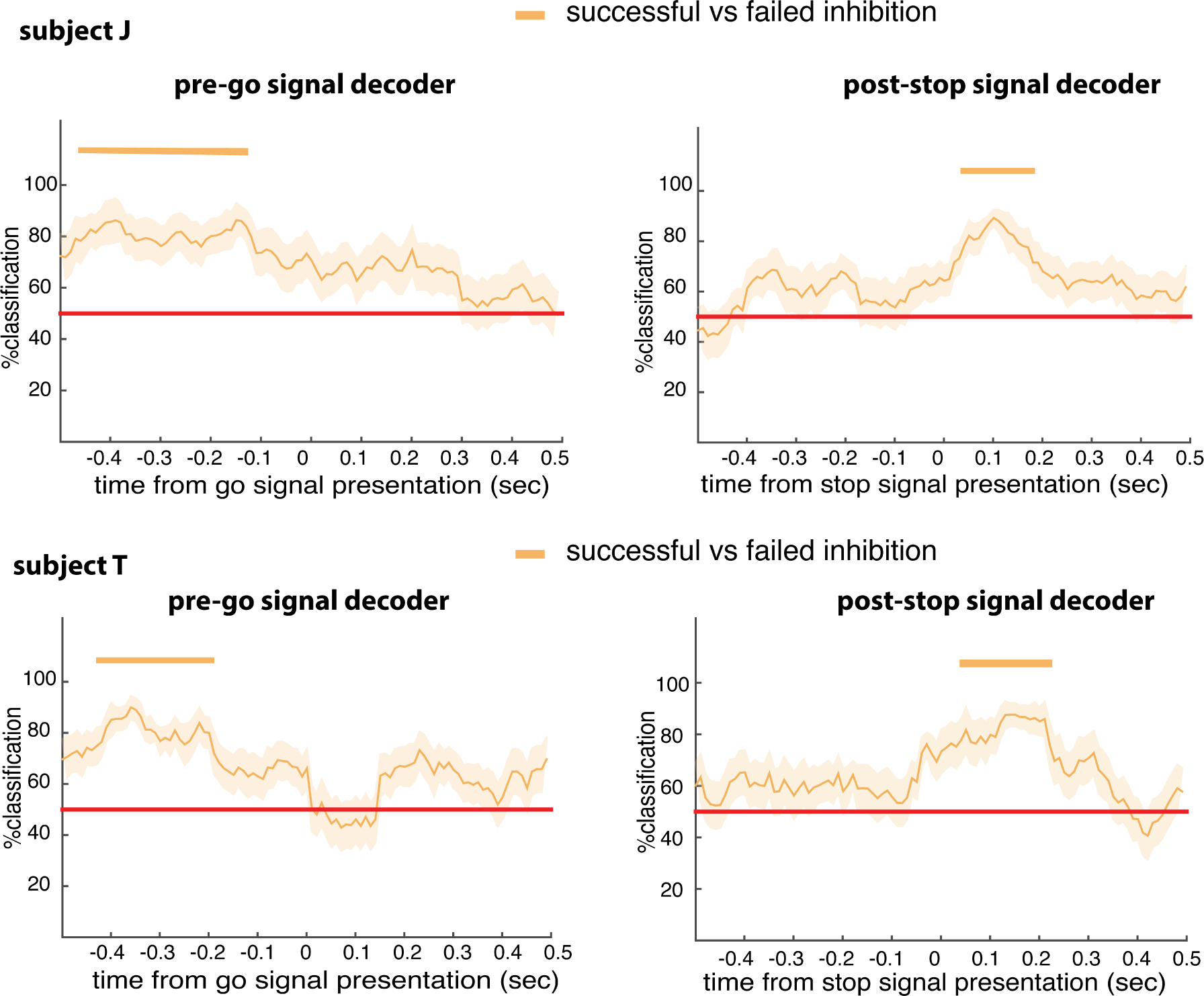
Post-stop and pre-go decoding for subjects J and T: The post stop signal decoder was able to classify success of an inhibition significantly above chance (see Methods for specific use of chi-square tests to quantify significance) in a time period ranging from 40 msec to 170 msec for subject J, and 40 to 220 msec for subject T, respectively after the stop signal (these times indicate the beginning of 100 msec boxcars, and chi-square tests were used for finding their significance with p < 0.05, see Methods). The first significant bin was therefore of window size 40 – 140 msec, that led to average cancellation time as 90 msec. It preceded the average stopping response by 50 msec in subject J, and by 30 msec in subject T, suggesting OFC’s responses may precede the stopping response. In Pre-go signal decoder, for subject J, high accuracy of decoding was found during the time periods 460 msec to 120 msec before the appearance of go signal. Likewise, it was 420 msec to 200 msec in subject T.

**Figure S2.**
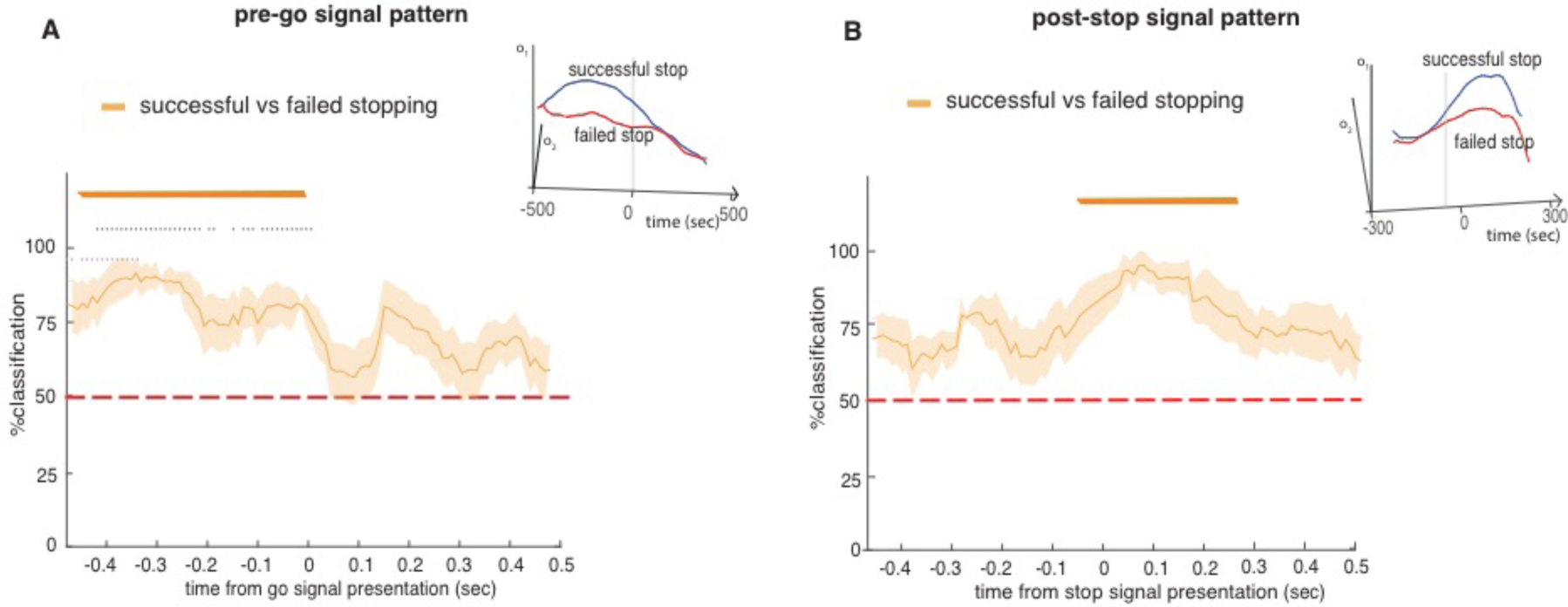
Post-stop and pre-go decoding for subjects J and T with zscored data (normalization): Both post-stop signal and pre-go signal decoder was able to classify success of stopping significantly above chance (see **Methods** for specific use of chi-square tests to quantify significance) before SSRT and go signal presentation, respectively, and chi-square tests were used for finding their significance with p < 0.05, see **Methods**). Results suggest that successful differentiation of stopping codes can be obtained irrespective of the normalization methods used in the study (In the manuscript, Normalization procedure was carried out by subtracting the mean firing during inter-trial interval (ITI) time period (baseline) and then by zscoring each neuron’s data, and the normalized data is used for decoder analysis).

